# Antibody response patterns to *Helicobacter pylori* infection in a rural Ugandan population cohort

**DOI:** 10.1101/2021.03.01.433355

**Authors:** Neneh Sallah, Alexander Hayes, Nana Osei-Tutu, Julia Butt, W. Thomas Johnston, Gershim Asiki, Tim Waterboer, Martin L. Hibberd, Robert Newton

## Abstract

**Background:** Helicobacter pylori (*H. pylori*) establishes life-long infection in humans in the absence of treatment and has been associated with a variety of gastrointestinal conditions including peptic ulcer and gastric cancer. Antibody responses to *H. pylori* antigens are found to be associated with disease risk, however, data from Africa are scarce.

**Methods:** To assess the seroprevalence of *H. pylori* and characterise antibody response patterns, we measured serum IgG antibody levels to 14 antigens among 7,211 individuals in a rural Ugandan population cohort. Multivariate-adjusted linear regression models were fitted to investigate the influence of age, sex, and co-infection on antibody seroreactivity levels.

**Results:** *H. pylori* seroprevalence was 95% in our study population, with 94% of individuals seropositive in childhood (<15 years). In *H. pylori* positive individuals, we found a markedly high seroprevalence (~99%) and antibody levels to the high-risk antigens CagA and VacA, in addition to Cagδ. HSV-2 co-infection was significantly associated with higher IgG levels of CagA and VacA (OR=1.10, 95% C. I=1.05-1.16). HIV infection was associated with lowered IgG levels to CagA (OR=0.86, 95% C.I.=0.80-0.93), and HPV infection was associated with increased IgG levels to VacA (OR=1.16, 95% C.I.=1.11-1.21).

**Conclusions:** *H. pylori* in this population is ubiquitous from childhood, with a high prevalence and high seroreactivity levels of high-risk antigens, suggesting chronic active inflammatory responses in individuals that are indicative of risk of disease. Further investigation is warranted to fully understand the relationship between host, immunogenicity, and clinical outcomes to better stratify by risk and improve treatment.

**Key Messages:**

- Antibody responses to *H. pylori* antigens are found to be associated with risk of gastric cancer, however, despite the high seroprevalence in African populations, data from Africa are scarce. This is the first study of antibody response patterns and their determinants from an African population.
- Our study shows a population where *H. pylori* is ubiquitous from childhood, and seroprevalence of virulent antigens is distinctively high suggesting an increased of disease compared to other populations.
- We observe inter-individual variation in virulent antibody responses partly influenced by co-infection.
- We highlight crucial insights into antibody-based biomarkers of disease risk and reinforce the need for population-based *H. pylori* screening and treatment programmes for gastric cancer control.

## Introduction

*Helicobacter pylori* (*H. pylori*), is a gram-negative bacterium that colonises the gastric epithelium of the human host and is typically acquired in childhood via intrafamilial contact[1]. Globally, 4.4 billion individuals are estimated to be infected with *H. pylori* with substantial geographic variation in prevalence ranging from ~20-50% in developed countries to ~80% in developing countries[2]. While the majority of infections are asymptomatic, in the absence of treatment, *H. pylori* can evade the host’s immune responses and subsequently persist throughout a person’s lifetime. Chronic *H. pylori* infection is the strongest risk factor for several gastrointestinal conditions including, peptic ulcer, chronic atrophic gastritis, and gastric cancers worldwide[3]. In 2018, *H. pylori* was responsible for 810,000 (37%) new cases of cancers caused by an infectious agent and caused ~90% of non-cardia gastric cancers[3]. The relative risk of developing gastric cancer with the *H. pylori* infection is 17[4]. In sub-Saharan Africa, the estimated population attributable fraction (PAF) of non-cardia gastric cancer is 92%[4]. Between the 1960s and 2008, Uganda experienced a 7-fold increase in gastric cancer incidence[5]. In 2008, The Uganda Cancer Working Group recognised that in areas of endemic *H. pylori* infection, *H. pylori* predicated the multistage process that lead to the development of gastric cancer[5]. They recommended that *H. pylori* eradication be utilised as a prevention strategy in areas with a high incidence of gastric cancer[5]. This recommendation is shared by the International Agency for Research on Cancer (IARC) working group that population-based *H. pylori* screening and treatment programmes are needed for gastric cancer control[6].

While the understanding of potential disease outcome following *H.pylori* infection is limited, it is found to be mediated by the complex interplay between bacterial virulence factors, environmental factors, lower socioeconomic status and host factors[7]. During the course of infection, chronic inflammatory changes occur as the bacterium adapts to its host and modulates the immune system. Depending on the bacterium’s protein expression pattern and the host’s adaptive immune response distinctive antibody response patterns may result and potentially reflect virulence of the infecting bacterium. As an example, antibody responses to particular *H. pylori* antigens: CagA (cytotoxin-associated antigen A), VacA (vacuolating cytotoxin), GroEL (chaperonin GroEL), HP1564 (hypothetical protein) and HcpC (conserved hypothetical secreted protein) have been identified as being associated with development of chronic atrophic gastritis and gastric cancers[8, 9]. A recent study characterised the dynamics of antibody responses to 15 antigens in a healthy German population-based cohort with a *H. pylori* seroprevalence of 48% and found multiple seropositivity and higher antibody levels associated with increasing age, suggestive of persistent infection and lifelong stimulation of immune responses[10]. Despite the high seroprevalence in African populations, there is a paucity of data, particularly from the continent compared to other populations[11, 12]. To better understand the relationship between host and pathogen in individuals the investigation of distinctive antibody patterns and factors that could potentially lead to disease following chronic infection is essential. Therefore, in this study, we assess the seroprevalence of *H. pylori* and antibody patterns against 14 antigens in a rural African population cohort and explore the factors associated with inter-individual variation in antibody response.

## Methods

### Sample selection and ethics

The General Population Cohort (GPC) is a population-based cohort in rural south-west Uganda consisting of 25 neighbouring villages inhabited mainly by farmers[13, 14]. The Baganda are the predominant tribal group constituting ~70% of the population, with a substantial number of migrants who settled from neighbouring Rwanda. Blood samples from 7,211 GPC study participants, representing 11 self-reported ethnolinguistic groups, were collected during census and medical survey sampling rounds conducted in the study area in 1992, 2000 and 2008, as described previously[13]. Serum was tested for infections and the remainder was stored at -80 degrees Celsius in freezers in Entebbe prior to further serological testing. Informed consent was obtained from all participants either in conjunction with parental/guardian consent for under 18-year olds with signature or a thumbprint if the individual was unable to write. The study was approved by the Uganda Virus Research Institutes, Research Ethics committee (UVRI-REC) (Ref. GC/127/10/10/25), the Uganda National Council for Science and Technology (UNCST) and the London School of Hygiene & Tropical Medicine (LSHTM) Ethics Committee (reference number 17686).

### Helicobacter pylori multiplex serology

We quantified mean fluorescent intensity (MFI) of IgG antibodies to 14 *H. pylori* antigens using multiplex serology on the Luminex® platform based on glutathione-S-transferase (GST) fusion capture immunosorbent assays combined with fluorescent bead technology as previously described[15]. The 14 recombinant affinity purified *H. pylori* proteins that were used as antigens were: CagA (cytotoxin-associated antigen A), GroEL (chaperonin GroEL), HP1564 (hypothetical protein protein HP1564), VacA (vacuolating cytotoxin), HP0231 (hypothetical protein HP0231), HP0305 (hypothetical protein HP0305), HpaA (neuraminyllactose-binding hemagglutinin homolog), Cagδ (cag pathogenicity island protein δ), HyuA (hydantoin utilization protein A), CagM (cag pathogenicity island protein M), Catalase, HcpC (conserved hypothetical secreted protein - paralogue HcpA induces IFNγ), NapA (neutrophil activating protein (bacterioferritin)) and UreA (urease alpha subunit). Antibody seropositivity was defined as reactivity greater than the antigen-specific cut-off. *H. pylori* seropositivity was defined as seropositivity to at least 4 antigens as described previously[10, 15].

### Statistical analysis of variability in antibody responses

Statistical significance of differences in continuous variables, i.e. antibody reactivities (MFI values) and multiple seropositivity (number of antigens recognized) were analysed with the Wilcoxon two sample signed rank sum test. Fisher’s Exact test was used to test for differences in dichotomous variables, i.e. *H. pylori* seropositivity and antibody seroprevalence. All tests were performed two-sided. Analyses were performed with R software version 3.4. P-values below 0.05 were considered statistically significant. Pairwise-correlation between antigens for seropositive individuals were tested using Spearman’s rank-based test in R. We also investigated factors influencing IgG antibody response levels to *H. pylori* antigens. Antibody levels were log_10_ transformed and multivariate linear regression models were fitted to examine variables predictive of antibody levels. The variables tested were: sex, age/age group (categorized as 0-14, 15-24, 25-44 or ≥45 years), census (sampling) year (1992, 2000 or 2008) and infection status for Epstein-Barr virus (EBV), Hepatitis B virus (HBV), Hepatitis C virus (HCV), Human-immunodeficiency virus (HIV), Human papillomavirus (HPV) high risk types 16 and 18, Herpes simplex virus type 2 (HSV-2), Human T-lymphotropic virus (HTLV-1), Kaposi’s sarcoma-associated herpesvirus (KSHV), and Merkel cell polyomavirus (MCV). A multiple testing *p-*value of <0.005 adjusted for all variables was used to determine statistical significance unless specified otherwise.

## Results

### Helicobacter pylori seroprevalence in the GPC

In this study, we analysed the sera of 7,211 individuals between the ages of 0-98 years (median=16, IQR=10-35) for antibody responses to 14 different *H. pylori* antigens: CagA, GroEL, HP1564, VacA, HP0231, HP0305, HpaA, Cagδ, HyuA, CagM, Catalase, HcpC, NapA, and UreA. We categorised 95% of individuals as seropositive for *H. pylori* infection based on reactivity to at least 4 antigens (Table 1). While *H. pylori* seroprevalence was similar for both sexes (males 95.5% vs females 94%), we found that in the 25-44 years age group seroprevalence was significantly higher in males (97.3%) compared to females (94.9%) (p=0.03) (Figure 1). We also found that seroprevalence was significantly higher (>2%) in the oldest age group (45+) compared to the youngest age group (0-14) for both males (p=0.02) and females (p=0.01) (Figure 1). Seroprevalence was also similarly high across census years (94-96%) (Table1). Seroreactivity to multiple antigens was high (median=8.7, IQR=7-11) with 3375 individuals (47%) testing seropositive for >9 antigens (Supplementary Figure 1.A). We found no significant differences in the number of antigens recognised across age groups or between sexes (P>0.05) (Supplementary Figure 1.B). We compared the seroprevalences of all antibodies to the 14 antigens in seropositive (HP+) vs seronegative (HP-) individuals and found that antibody seroprevalences were significantly higher in HP+ compared to HP- sera for all antigens (p<0.0001) (Table2). In the 6,834 *H. pylori* seropositive individuals, antibodies to CagA, VacA and Cagδ were the highest with ~99% seroprevalence and antibodies to NapA and HcpC had the lowest seroprevalence being less than 50% (Table2).

**Table 1.**
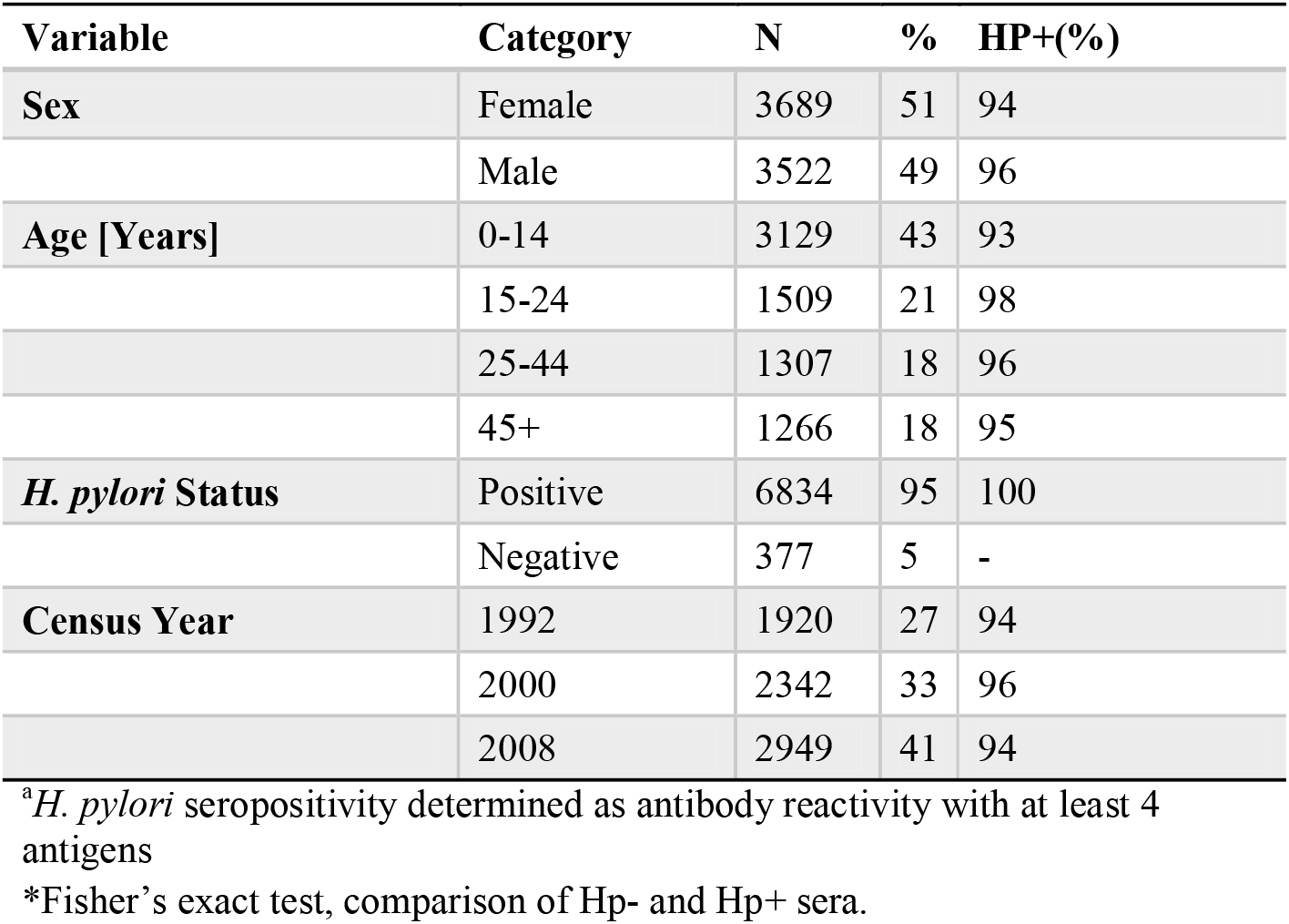
Characteristics of samples in the GPC (n=7211)

**Table 2.**
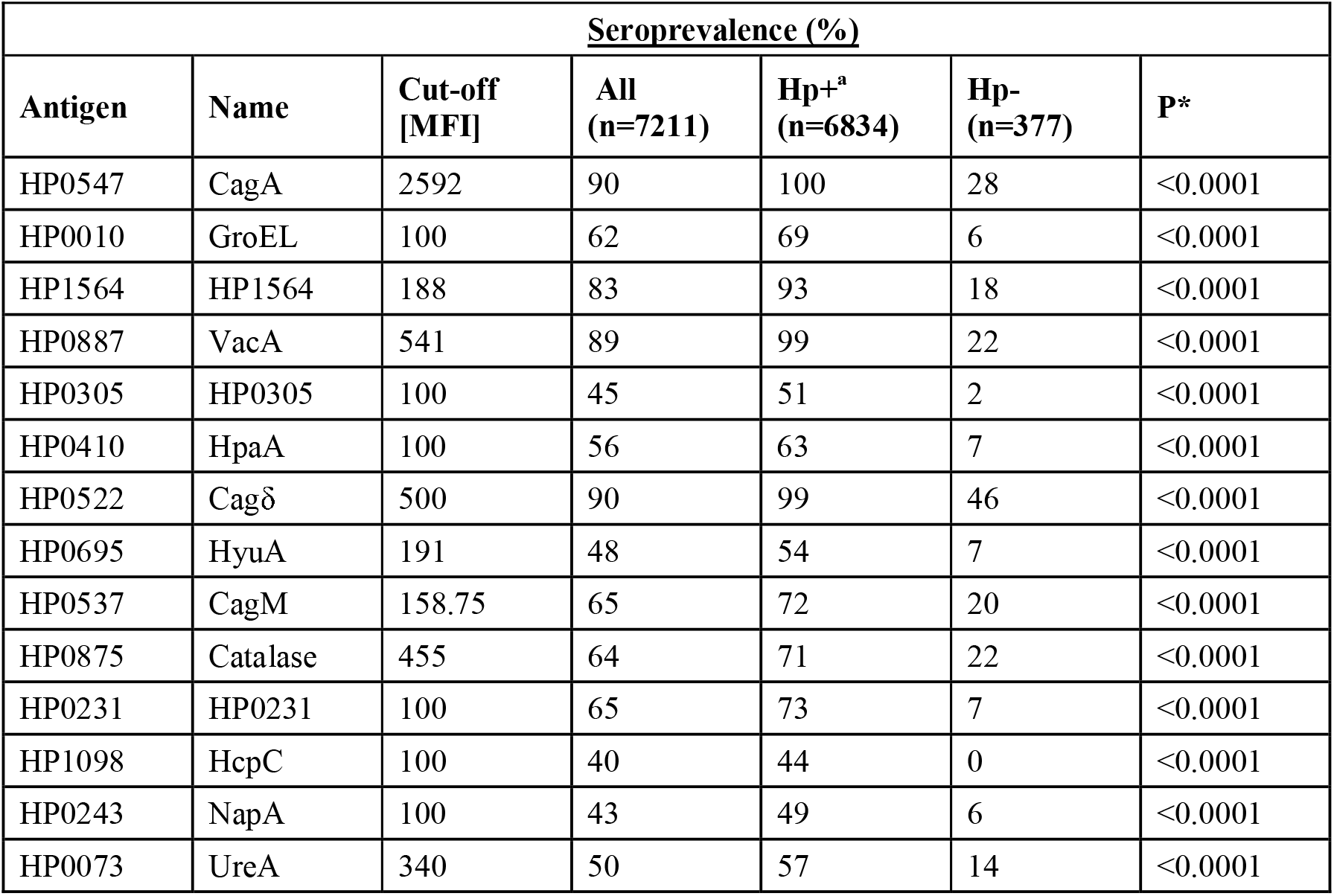
Seroprevalence of antibodies to *H. pylori* antigens by HP serostatus.

**Figure 1.**
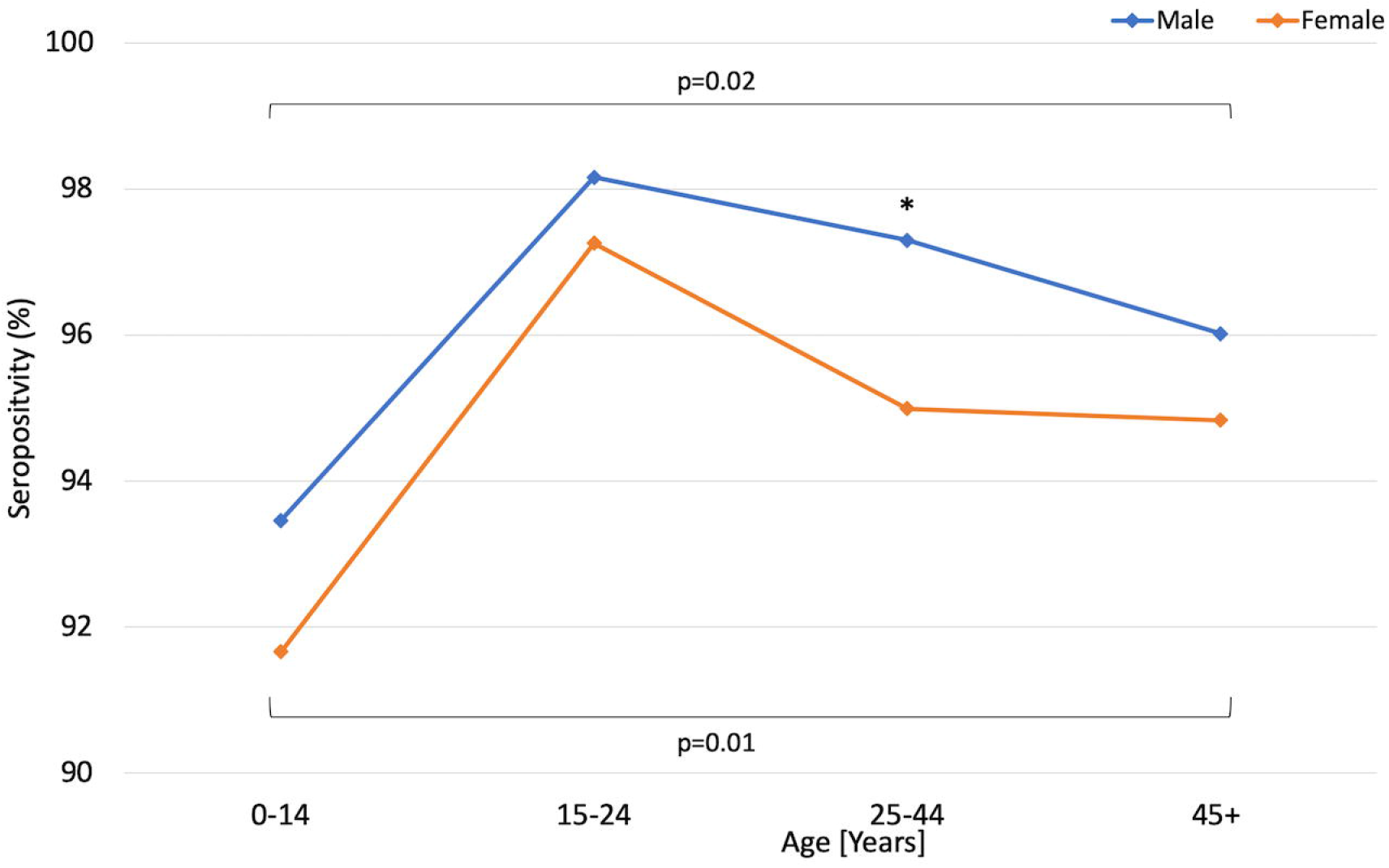
*H. pylori* seropositivity by gender and age. Star indicates statistically significant difference between genders in age 25-44, p=0.03 (Fisher’s exact test). Horizontal brackets indicate statistically significant differences between the youngest (0-14) and oldest (45+) age groups for both genders: Males: p=0.02, females: p= 0.01 (Wilcoxon two sample signed rank sum test).

### Helicobacter pylori antibody response levels

Using the multiplex flow immunoassay, we quantified the antibody levels of response to the 14 different *H. pylori* antigens. In *H.pylori* positive and antigen-specific antibody positive individuals, we observed variation in antibody responses with the highest IgG levels observed for CagA (max= 66,614, median=13,793, IQR=10,138-18,298) followed by VacA (max=28,722, median=2,809, IQR=1641-4600), HP1564 (max=22,526, median:2776, IQR=1341-4640) and Cagδ (max=15,044, median:2,206, IQR=1,293-3,609), all other antibody responses had medians below 2000 with HpaA having the lowest median IgG level (max= 10,783, median:335, IQR:182-751) (Figure 2.). There was a modest correlation between all antigens (rho=-0.01-0.51) (Supplementary Figure S2), with the strongest correlation being observed for HP0305 and HP1564 (rho=0.51) (Supplementary Figure S2).

**Figure. 2.**
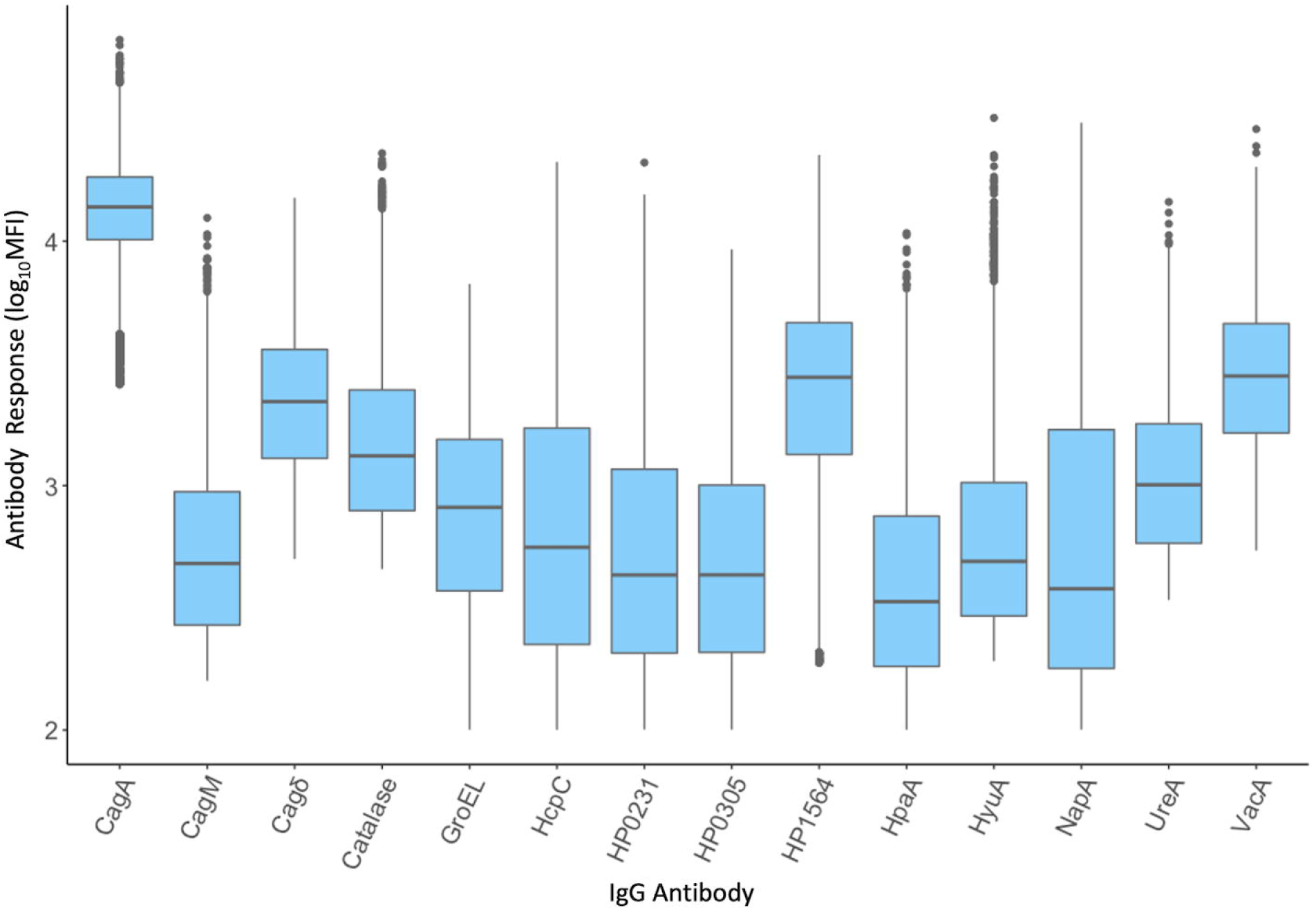
Distribution of antibody responses (log_10_MFI) in H. pylori positive and antigen positive individuals. Horizontal lines denote median and vertical lines denote interquartile range (IQR).

We investigated the association between intrinsic factors, age, sex and sampling year to inter-individual variation in antibody response levels in seropositive individuals using a multivariable linear regression model for each antibody (Table 3). We found that higher age groups in comparison to the youngest age group (0-14) were significantly associated with decreased antibody levels to CagA, HP1564, VacA and HP0231 (ORs between 0.84-99) (Table 3). In contrast, higher age groups were significantly associated with increased antibody levels to GroEL, Cagδ, HyuA, NapA, HcpC and UreA (ORs between 1.01-1.21) in comparison to the youngest age group (Table 3). Being female was significantly associated (p<0.0125) with lower antibody responses to HP1564, HP0305, Cagδ, HyuA and Catalase (ORs between 0.91-99) (Table 3) compared to being male.

**Table 3.** Predictors of antibody response levels to *H. pylori* antigens in seropositive individuals.

|  |  |  |  |  |  |  |
| --- | --- | --- | --- | --- | --- | --- |
| Variable | Age Group: Ref =0-14 years |  |  | Sex:<br>Ref=Male | Census Year: Ref=1990 |  |
| Antigen | 15-24 | 25-44 | 45+ | Female | 2000 | 2008 |
| **CagA** |  |  |  |  |  |  |
| OR (95% C.I) | 0.98 (0.96 - 0.99) | 0.96 (0.95 - 0.98) | 0.91 (0.90 - 0.93) | 1.00 (0.99 - 1.01) | 1.03 (1.01 - 1.04) | 1.04 (1.03 - 1.06) |
| p-value | 0.002 | <0.001 | <0.001 | 0.540 | <0.001 | <0.001 |
| **GroEL** |  |  |  |  |  |  |
| OR (95% C.I) | 1.08 (1.04 - 1.11) | 1.06 (1.03 - 1.1) | 1.10 (1.06 - 1.14) | 0.98 (0.96 - 1.01) | 1.04 (1.01 - 1.07) | 1.11 (1.07 - 1.14) |
| p-value | <0.001 | 0.001 | <0.001 | 0.153 | 0.019 | <0.001 |
| **HP1564** |  |  |  |  |  |  |
| OR (95% C.I) | 0.95 (0.92 - 0.97) | 0.89 (0.87 - 0.92) | 0.86 (0.84 - 0.89) | 0.93 (0.91 - 0.95) | 1.02 (0.99 - 1.05) | 0.99 (0.97 - 1.02) |
| p-value | <0.001 | <0.001 | <0.001 | <0.001 | 0.146 | 0.699 |
| **VacA** |  |  |  |  |  |  |
| OR (95% C.I) | 1.01 (0.99 - 1.03) | 0.99 (0.97 - 1.01) | 0.97 (0.94 - 0.99) | 0.98 (0.97 - 1.00) | 1.03 (1.01 - 1.05) | 1 (0.98 - 1.02) |
| p-value | 0.224 | 0.399 | 0.002 | 0.019 | 0.004 | 0.701 |
| **HP0305** |  |  |  |  |  |  |
| OR (95% C.I) | 0.94 (0.91 - 0.98) | 0.94 (0.90 - 0.98) | 0.95 (0.91 - 1.00) | 0.94 (0.91 - 0.97) | 1.01 (0.97 - 1.05) | 1.05 (1.01 - 1.09) |
| p-value | 0.004 | 0.004 | 0.041 | <0.001 | 0.523 | 0.013 |
| **HpaA** |  |  |  |  |  |  |
| OR (95% C.I) | 1.03 (0.99 - 1.06) | 0.98 (0.94 - 1.02) | 1.02 (0.98 - 1.06) | 0.97 (0.95 - 1.00) | 1.04 (1.00 - 1.07) | 1.03 (0.99 - 1.06) |
| p-value | 0.106 | 0.290 | 0.257 | 0.023 | 0.026 | 0.102 |
| **CagD** |  |  |  |  |  |  |
| OR (95% C.I) | 1.03 (1.01 - 1.05) | 1.04 (1.02 - 1.06) | 1.03 (1.01 - 1.05) | 0.95 (0.93 - 0.96) | 1.01 (1.00 - 1.03) | 0.98 (0.97 - 1.00) |
| p-value | 0.002 | 0.001 | 0.005 | <0.001 | 0.143 | 0.112 |
| **HyuA** |  |  |  |  |  |  |
| OR (95% C.I) | 1.02 (0.98 - 1.06) | 1.12 (1.07 - 1.16) | 1.15 (1.10 - 1.19) | 0.96 (0.93 - 0.99) | 1.01 (0.97 - 1.05) | 1.02 (0.98 - 1.06) |
| p-value | 0.254 | <0.001 | <0.001 | 0.003 | 0.613 | 0.325 |
| **CagM** |  |  |  |  |  |  |
| OR (95% C.I) | 1.01 (0.98 - 1.04) | 0.99 (0.96 - 1.03) | 0.99 (0.96 - 1.03) | 0.98 (0.96 - 1.01) | 1.01 (0.98 - 1.04) | 1.03 (1.00 - 1.06) |
| p-value | 0.408 | 0.746 | 0.686 | 0.151 | 0.475 | 0.022 |
| **Catalase** |  |  |  |  |  |  |
| OR (95% C.I) | 0.98 (0.96 - 1.01) | 0.98 (0.95 - 1.01) | 0.99 (0.96 - 1.02) | 0.97 (0.95 - 0.99) | 1.01 (0.99 - 1.04) | 1.03 (1.01 - 1.06) |
| p-value | 0.169 | 0.173 | 0.485 | 0.001 | 0.319 | 0.008 |
| **HP0231** |  |  |  |  |  |  |
| OR (95% C.I) | 0.96 (0.93 - 1.00) | 0.93 (0.89 - 0.97) | 1.00 (0.96 - 1.04) | 0.97 (0.94 - 1.00) | 1.01 (0.98 - 1.05) | 1.06 (1.03 - 1.1) |
| p-value | 0.028 | 0.001 | 0.838 | 0.023 | 0.515 | <0.001 |
| **HcpC** |  |  |  |  |  |  |
| OR (95% C.I) | 1.00 (0.95 - 1.05) | 1.06 (1.00 - 1.13) | 1.13 (1.07 - 1.20) | 0.96 (0.92 - 1.00) | 1.03 (0.98 - 1.09) | 1.04 (0.99 - 1.09) |
| p-value | 0.924 | 0.049 | <0.001 | 0.044 | 0.219 | 0.137 |
| **NapA** |  |  |  |  |  |  |
| OR (95% C.I) | 1.13 (1.07 - 1.20) | 1.14 (1.07 - 1.21) | 1.13 (1.06 - 1.20) | 1.00 (0.95 - 1.04) | 0.98 (0.92 - 1.03) | 1.00 (0.95 - 1.06) |
| p-value | <0.001 | <0.001 | <0.001 | 0.813 | 0.389 | 0.879 |
| **UreA** |  |  |  |  |  |  |
| OR (95% C.I) | 1.01 (0.98 - 1.04) | 1.02 (0.99 - 1.05) | 1.07 (1.04 - 1.10) | 0.97 (0.95 - 0.99) | 1.00 (0.97 - 1.03) | 1.02 (0.99 - 1.05) |
| p-value | 0.395 | 0.159 | <0.001 | 0.012 | 0.864 | 0.120 |

### The influence of co-infection on Helicobacter pylori antibody responses

We then investigated the seroprevalence of co-infection in 1,735 individuals (aged 1-91 years, median= 31) that had a record of infection serostatus for nine additional pathogens. We found that in 1,679 (97%) *H.pylori* positive individuals there was a high burden of co-infection, particularly with Epstein-Barr Virus (EBV) and Kaposi’s Sarcoma-associated herpesvirus (KSHV) (93%) (Figure 3.A.) followed by Herpes-simplex virus-2 (HSV-2) (67%), Human Papilloma Virus (HPV) (35%) and Hepatitis B Virus (HBV) (13%). The remaining four pathogens were <10% with Human T lymphotropic virus-1 (HTLV-1) having the lowest seroprevalence (~1%) (Figure 3.A.). Most individuals (~70%) were co-infected with at least four pathogens (Figure 3.B).

**Figure 3.**
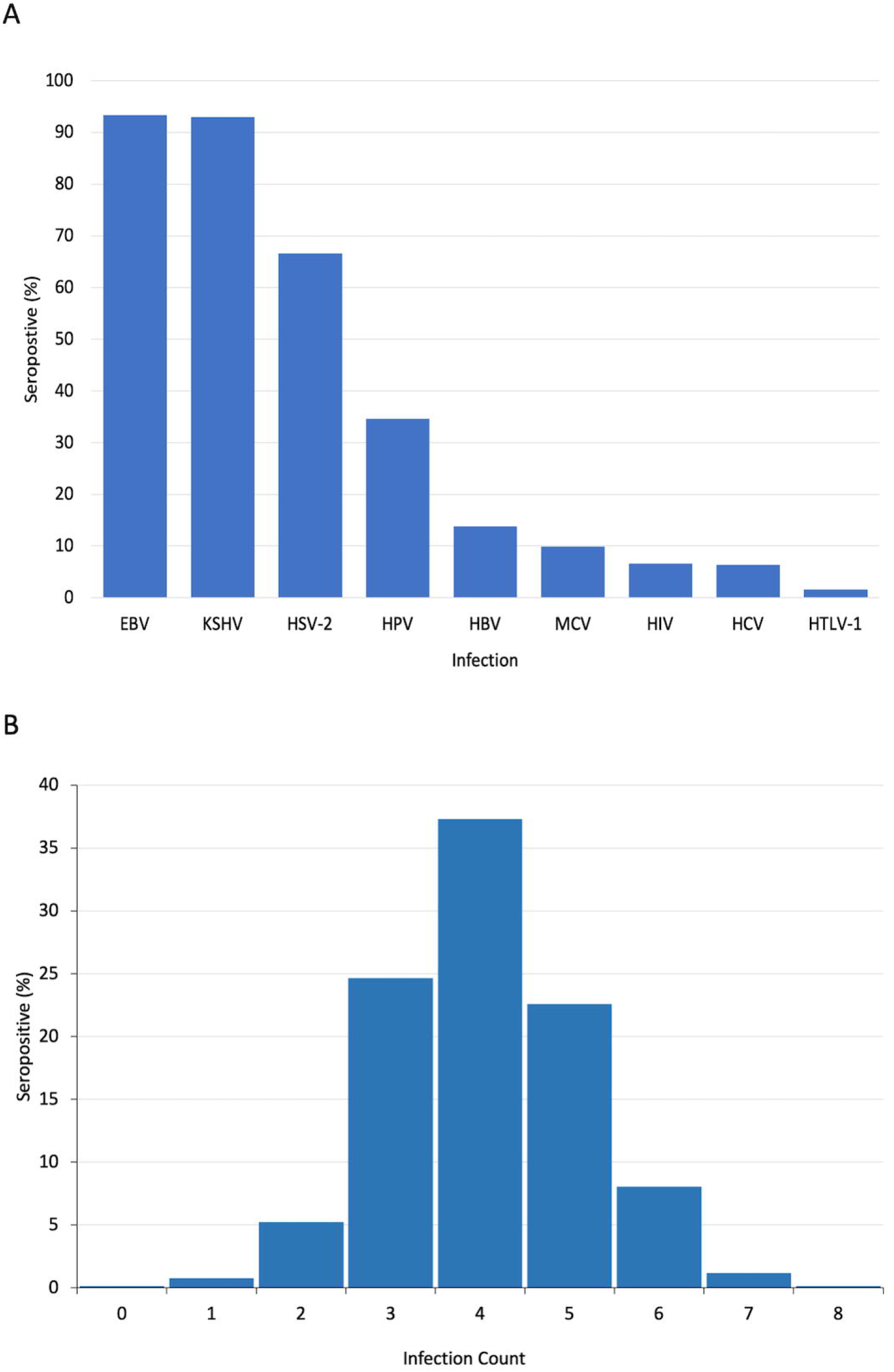
A. Seroprevalence of co-infection in *H. pylori* seropositive individuals (n=1679). EBV: Epstein Barr Virus, KSHV: Kaposi’s Sarcoma-associated Herpesvirus, HSV-2: Herpes Simplex Virus-2, HBV: Hepatitis B virus, HPV: Human papilloma virus, MCV: Merkel Cell polyoma virus, HIV: Human Immunodeficiency virus, HCV: Hepatitis C virus and HTLV-1: Human T-cell Lymphotrophic virus. **B. Burden of infection**. Infection count represents the number of infections the individuals tested positive for.

We then investigated the influence of co-infection on inter-individual variation in antibody response levels to the five virulent *H. pylori* antigens, CagA, VacA, GroEL, HcpC and HP1564, in *H.pylori* seropositive individuals using multivariable linear regression models adjusting for age, sex, sampling year and infection status. Of the infections tested (Figure 3. A), HIV co-infection resulted in significantly lowered antibody responses to CagA, HcpC, and HP1564 (OR= 0.55-0.93) (Table 4). HSV-2 co-infection was associated with moderately higher levels of CagA and VacA (OR= 1.05-1.16) but no other antibodies (Table 4). HPV co-infection was significantly associated with higher antibody levels (OR= 1.11-1.45) to VacA and GroEL antigens (Table 4). None of the other co-infections had significant effects on variation for any antibody response (p>0.004) (Table 4).

**Table 4.**
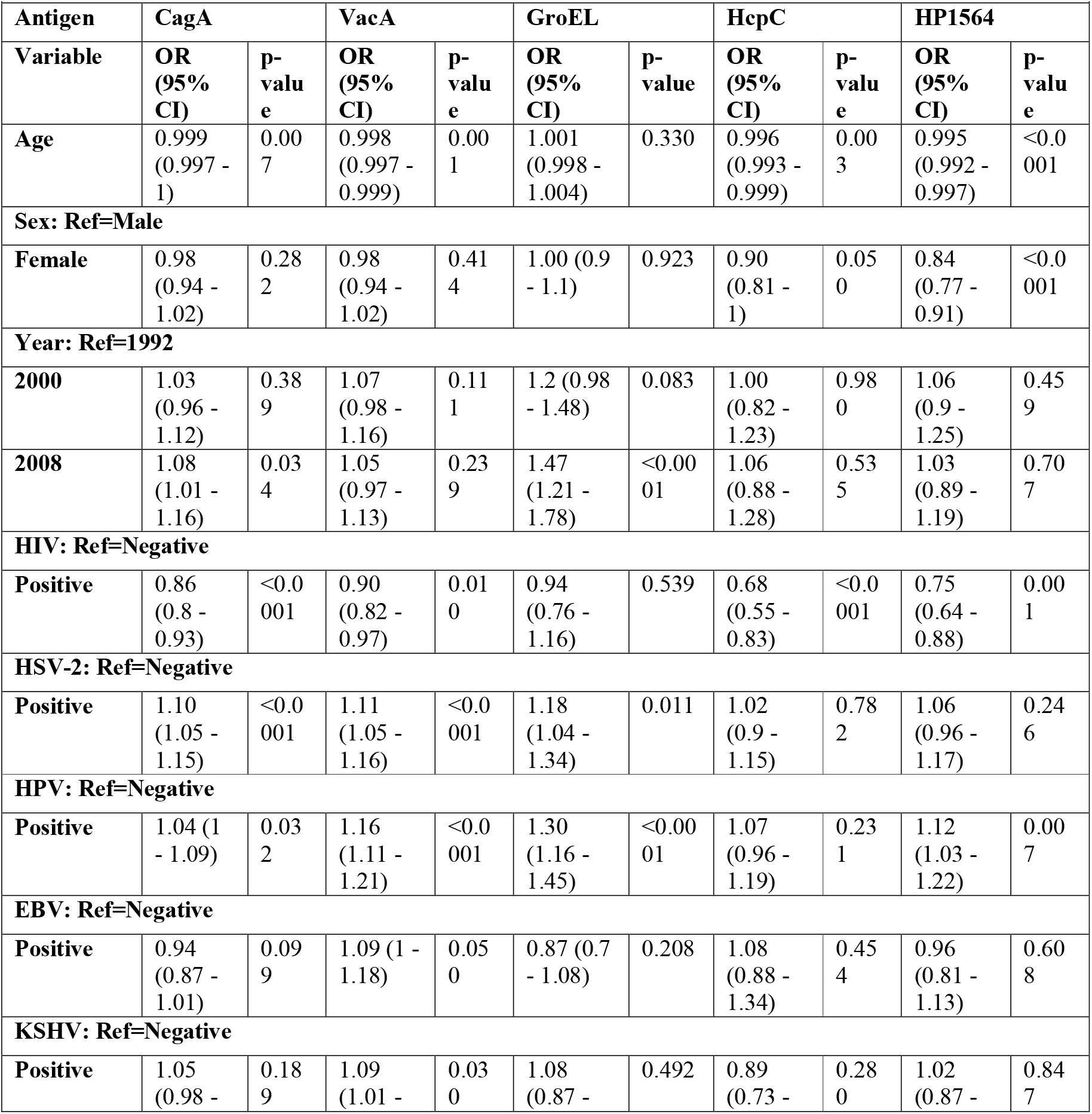
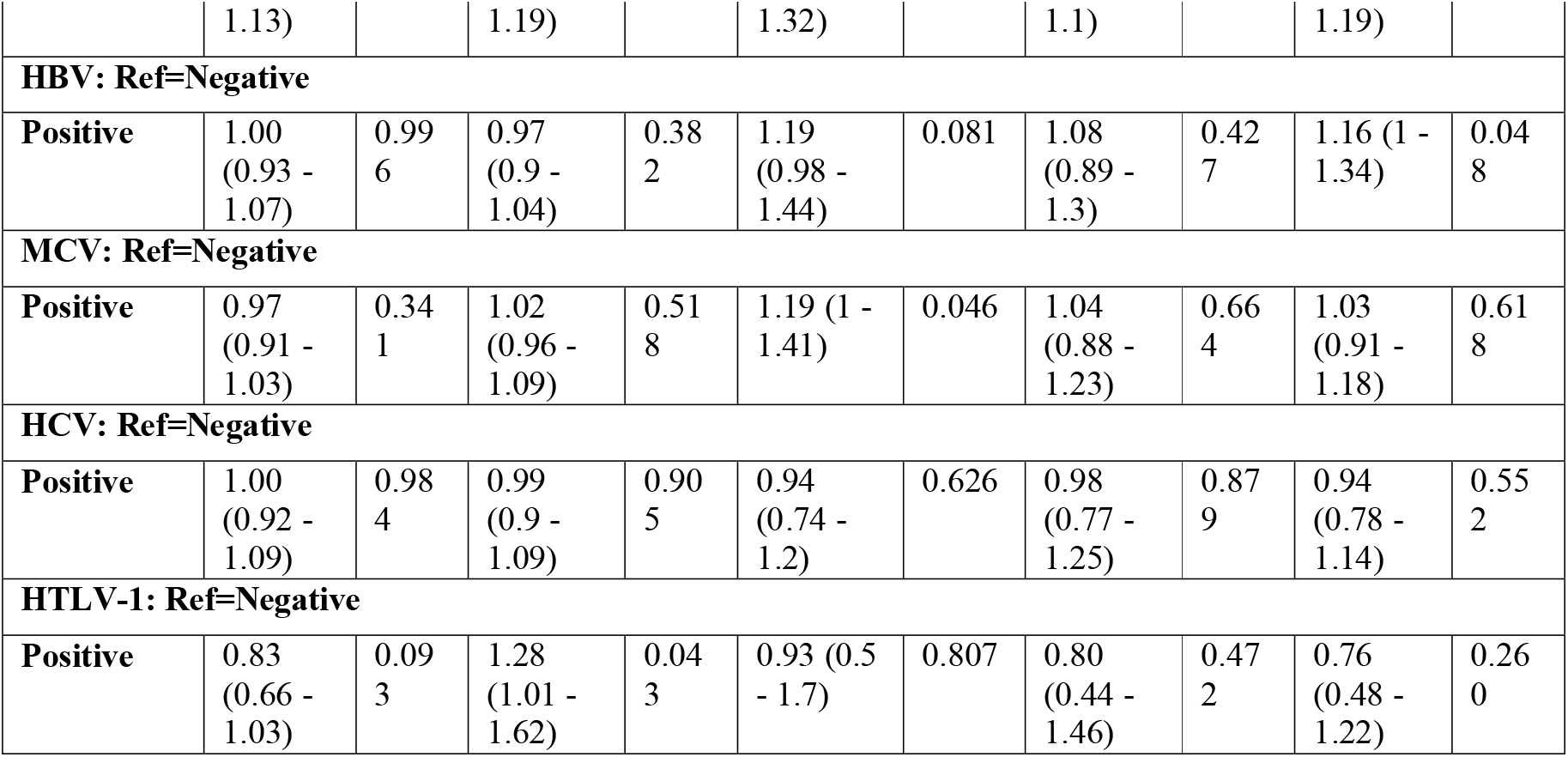
The influence of co-infection on antibody response levels to virulent *H. pylori* antigens (N=1679)

## Discussion

Data on *Helicobacter pylori* antibody response patterns and their determinants are limited, with most studies having been conducted in Western and non-African populations, there is a paucity of data particularly from Africa. Here, we present the first population-based study to characterise *H. pylori* antibody response patterns to 14 antigens and their determinants in ~7000 individuals from an African population. The presence of antibodies representing host immune response is commonly used as diagnostic markers for the stage of infection [16] and for *H.pylori* they have been found to be associated with disease risk[8, 9]. In the GPC, prevalence estimates of *H. pylori* of ~94% are higher than previous findings in Uganda and more broadly in developing countries[17-21]. Here, we see a population with a high seroprevalence of >90% from childhood, much higher than previously observed in Uganda of 44%; a seroprevalence higher than in other African countries, with studies reporting estimates in children ranging from 14% in Ghana to 73% in Kenya[11, 12, 17, 22]. We observed slightly higher seroprevalence in the highest age group compared to the lowest age groups (p<0.05), consistent with previous findings in other populations. In comparison to a previous study of German individuals[10] that assessed the presence of the 14 antigens tested here, we observed a 2-fold increase in seropositivity to *H. pylori* (95% in the GPC vs 48% in Germans). In *H. pylori* seropositive individuals, we observed >90% seropositivity to the following antibodies: CagA, HP1564, VacA and Cagδ, which was much higher than reported in previous studies. The highest difference in seroprevalence (2.7-fold) was observed for CagA (90% vs 33% Germans). For the following antigens: Cagδ, HP0231, HpaA, CagM and Catalase we also observed a >2-fold difference in seroprevalence with higher estimates in Ugandans compared to Germans. The lowest difference in seroprevalence was observed for HcpC, 1.17-fold higher in Ugandans compared to Germans. Another study in adults from high-risk gastric cancer areas of Latin America also reported lower antibody seroprevalences compared to our study, including for CagA (74%) and VacA (71%) despite an overall *H. pylori* seroprevalence of 85%[23]. Differences in seropositivity have also been reported in a low-income population in the U.S.A that had a higher *H. pylori* seropositivity 89% compared to the general US population (~30%), thought to be driven by lower socio-economic status[24]. They also observed higher seroprevalence in and higher in African Americans 79% compared to 69% in whites in addition to higher antibody seroprevalence for 12 out of the 14 antigens tested here, particularly for CagA, with a 2.6-fold increase in African Americans (68%) compared to white Americans (26%)[24]. This could be suggestive of higher seroprevalence in individuals with African ancestry. While seroprevalence estimates of CagA in Africa are scarce, the ubiquity of CagA in the GPC mirrors estimates in East Asian countries where the risk of gastric cancers due to *H. pylori* are higher than the rest of the world[3, 25]. CagA and VacA are the most well studied virulence factors of *H. pylori* and are essential for bacterial persistence in the stomach in addition to possessing oncogenic potential. CagA-host interaction induce a state of chronic inflammation which worsens over time and has been associated with an increased risk of gastric cancers, peptic ulcers, and other complications[26-28]. Studies that have evaluated antibody response levels to CagA and VacA report high antibody levels are associated with gastric mucosal inflammation, the grade of histological gastritis, and gastric cancer risk[29, 30]. It is interesting to observe such high seropositivity and immunogenicity to the high-risk serotypes, CagA and VacA in the GPC compared to non-African populations. A recent study in individuals in Southeast USA also found significantly higher levels of antibodies to CagA and VacA in African Americans compared to whites[31] that was likely driven by differences in ancestry.

Differences in antibody responses compared to other populations could occur as a result of multiple factors particularly, host genetic and environmental variation. In the GPC, a high burden of co-infection exists, with most individuals having up to four infections (Figure 3), therefore we sought to investigate whether inter-individual variation in antibody responses to the five virulent antigens CagA, VacA, HP1564, HcpC and GroEL was influenced by co-infection with pathogens previously reported to have oncogenic potential[3]. Out of the infections tested, only HIV, HSV-2 and HPV (high risk genotypes 16 or 18) co-infection influenced antibody levels. We found that while HIV infection was associated with decreased antibody responses (OR=0.53-0.99), co-infection with HSV-2 or HPV were significantly associated with moderate increases in antibody responses (OR=1.05-1.45) (Table 4). Co-infections can cause an imbalance in host immune system, likely modulating shared inflammatory pathways and thus, resulting in antibody variation[32-35]. However, as effect sizes in this study are small, they only partially explain inter-individual differences in antibody patterns.

In summary, we characterised antibody response patterns to 14 *H. pylori* antigens in ~7,000 individuals from a rural Ugandan population cohort and identified factors associated with inter-individual variation in response. A major finding of this study is that *H. pylori* seropositivity is ubiquitous from childhood (94%), and higher than previous reports in Uganda and other countries which could be due to lower-socioeconomic status in this cohort compared to urban areas. We also found that antigen specific seropositivity to the 14 antigens tested here, particularly for CagA, are higher than for non-African populations. Interindividual variation in antibody responses was partly explained by HIV, HSV-2 and HPV co-infections, suggesting other factors not examined here such as host genetics might be important. These insights are useful to better stratify individuals at risk of developing adverse outcomes following infection, as this population sustains such high levels of *H. pylori*, further investigation will be essential to inform appropriate therapeutic interventions to prevent disease in those at risk. Future studies would need to assess the relationship between antibody responses and clinical outcomes, and also investigate the relationship between host genetic variation and antibody variation to fully understand disease risk and population specific differences that are not explained by environmental variation.

## Acknowledgements

We thank all study participants who contributed to this study and acknowledge the support of the MRC/UVRI & LSHTM Uganda Research Unit

## Author Contributions

N.S, R.N., T.W. & M.L.H. designed the study. N.S & R.N wrote the manuscript. J.B & T.W performed phenotype assay development and validation. W.T.J, carried out curation of the cohort data and managed the database. N.S, A.H & N.O carried out all statistical analyses and data visualisation. N.S, R.N. & J.B. interpreted the results. G.A was the programme leader for the GPC. All authors commented on the interpretation of the results, reviewed and approved the final manuscript.

