## Supplemental files for "Antibody response patterns to *Helicobacter pylori* infection in a rural Ugandan population cohort"

Sallah et al.

**Supplementary Figures**

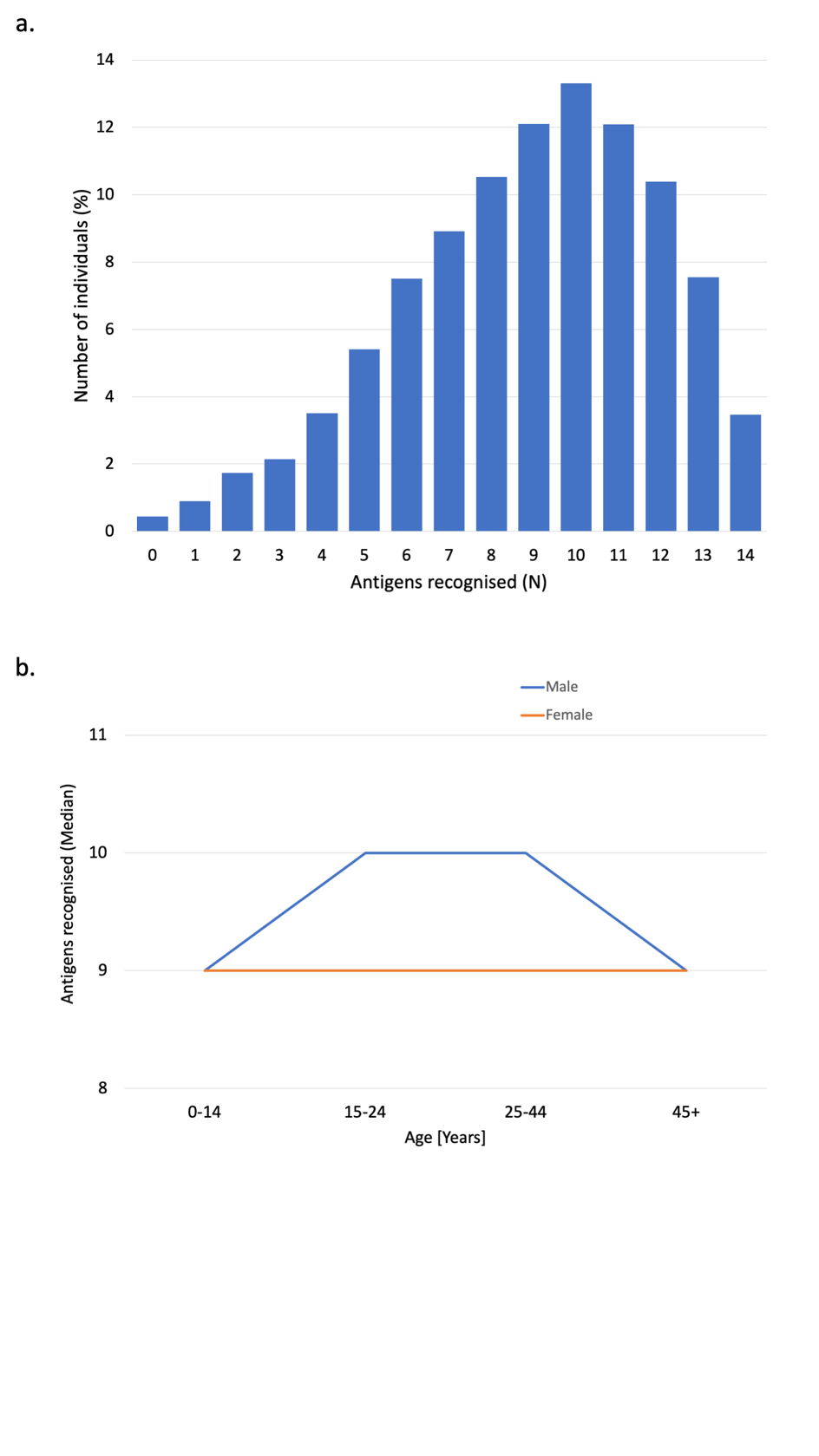

**Supplementary Figure 1.** **A.** Number of antigens recognised in 7211 individuals. **B.** The median number of antigens recognised within gender across age groups. Fisher’s exact test found no significant difference between genders within age groups. Wilcoxon-2-sample-signed test found no significant difference between the youngest and oldest age group by gender

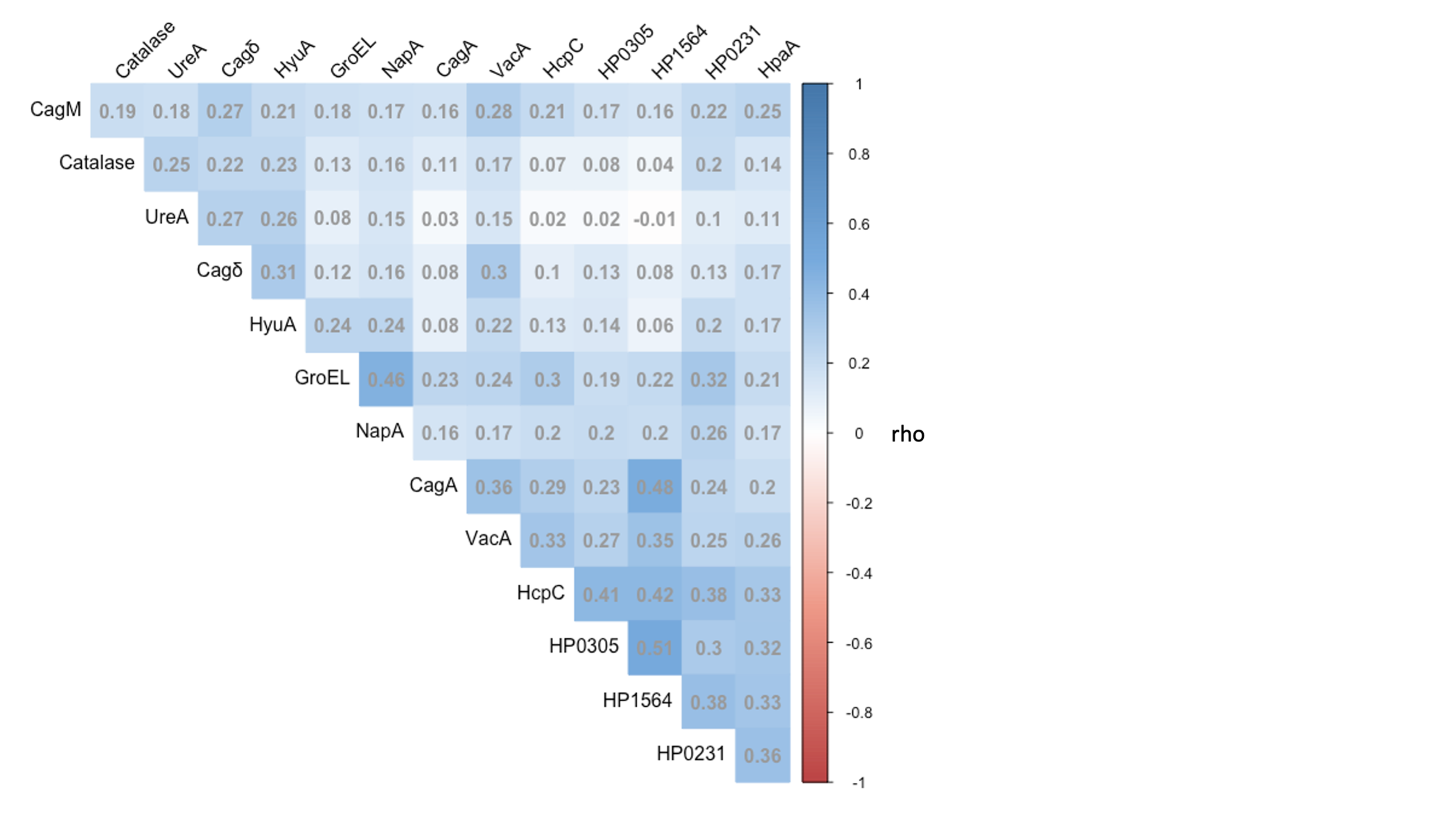

**Supplementary Figure 2.** Correlation matrix of antibody responses (MFI) for *H. pylori* antigens. Positive correlations are in blue and intensity is proportional to the correlation coefficients (rho) labelled in the squares and indicated on the right-hand side of the correlogram. All tests meet Spearman’s significance threshold of p<0.05.
